# IL-12R signaling promotes type 1 regulatory T cell specialization by sustaining T-bet

**DOI:** 10.64898/2026.01.10.698646

**Authors:** Anna Estrada Brull, Ylva M. Carlen, Laura Revert Rubio, Pascale Zwicky, Sarah Mundt, Burkhard Becher, Nicole Joller

## Abstract

Elucidating how regulatory T (Treg) cells specialize and acquire specific suppressive capacities is key for understanding immune regulation in autoimmunity and cancer. Although IL-12 is a key driver of Th1 differentiation, its role in Treg cells remains poorly defined. Here, we investigated whether IL-12R signaling contributes to Treg cell specialization during Th1 immune responses. We show that IL-12Rβ2 KO Treg cells display reduced type 1 specialization following LCMV infection, resulting in impaired suppressive capacity. Mechanistically, IFN-γ induces a first peak of T-bet, whereas IL-12 acts downstream to sustain T-bet through a positive feedback loop. The decreased suppressive function of IL-12Rβ2 KO Treg cells alters effector T cell responses during both acute and chronic infections. Importantly, human Treg cells also respond to IL-12, promoting enhanced type 1 specialization. Together, these findings establish IL-12 as a critical regulator of type 1 Treg specialization and highlight the therapeutic potential of targeting this pathway.

**Teaser:** IL-12R signaling in Treg cells enables sustained T-bet expression, programing type 1 Tregs for suppression of Th1 responses.

## Introduction

Regulatory T cells (Treg cells) are key players in the maintenance of homeostasis and establishment of immune tolerance. Treg cells are CD4^+^ T cells characterized by the expression of the transcription factor Foxp3, which confers them with suppressive potential(*1*). During infections, Treg cells play both protective and detrimental roles: they can limit immunopathology and promote tissue repair(*2*), but at the cost of impeding pathogen clearance(*3*, *4*).

Infection settings also reveal the capacity of Treg cells to undergo functional specialization. In these contexts, Treg cells can adopt features of conventional CD4^+^ Foxp3^−^ T cell subsets (Tconv), mirroring their transcription factor and signature marker expression. While naive Tconvs differentiate to Th1 cells through the upregulation of the transcription factor T-bet(*5*), Treg cells acquire T-bet and a Th1-like phenotype (from now on referred as type 1 Treg cell specialization) during viral infections. These T-bet^+^ Treg cells have been shown to be essential for control of exacerbated Th1 responses(*6*, *7*).

T-bet expression in Th1 cells is predominantly driven by IL-12(*8*, *9*). IL-12 is a potent proinflammatory cytokine described in the context of type 1 immune responses. It is sensed through a heterodimeric receptor composed of IL-12Rβ1 (common subunit with other cytokine receptors, such as IL-23R) and IL-12Rβ2 (specific for IL-12 and IL-35). While IL-12Rβ1 is constitutively expressed, IL-12Rβ2 expression in Tconv cells is induced after infection through the interferon-γ (IFN-γ) – T-bet pathway and, subsequently, IL-12 itself(*10*, *11*). IL-12R signaling in Th1 cells plays a central role for T-bet expression after the first 72h post-infection and results in a higher Th1 polarization and IFN-γ production in recall responses(*12*).

For Treg cells, IFN-γ and to a lesser degree IL-27 have been described to be the main drivers of T-bet expression(*6*, *13*), whereas a role for IL-12 in T-bet induction in Treg cells remains to be addressed. Prior reports suggested that the lack of IL-12R signaling in Treg cells could serve as a protective mechanism against an ineffective effector-like phenotype(*14*). However, other groups, including our own, observed the upregulation of IL-12Rβ2 expression in Treg cells after LCMV infection(*15*, *16*). Hence, we aimed to understand whether the upregulation of a functional IL-12R is involved in Treg cell specialization after type 1 immune challenge.

Here, we made use of Treg cell-specific *Il12rb2* conditional Knock-Out (cKO) mice and the murine model of LCMV to show that IL-12R signaling contributes to type 1 Treg cell specialization. IL-12Rβ2 KO Treg cells display impaired acquisition of T-bet, CXCR3 and type 1-specific co-inhibitory receptor expression upon LCMV infection, resulting in reduced suppressive function. Mechanistically, IL-12 acts after the IFN-γ-mediated T-bet induction to sustain T-bet expression throughout the course of the infection, whereas the role of IFN-γ appears to be transient. The decreased suppressive capacity of IL-12Rβ2 KO Treg cells allows for effector T cells to proliferate more strongly and produce higher amounts of cytotoxic molecules during acute responses, and modulates the establishment of exhaustion in CD8^+^ T cells during chronic infections. Finally, we show that human Treg cells also respond to IL-12 *in vitro* and acquire a stronger type 1 specialization than in response to IFN-γ.

## Results

### *Il12rb2* is upregulated in Treg cells after LCMV infection

To determine the dynamics of IL-12Rβ2 expression in Treg cells in a Th1 dominated immune response, we infected Foxp3 GFP-KI mice with LCMV WE, which causes an acute, systemic Th1 response. Treg cells and CD44^hi^ Tconv cells were FACS-sorted and their *Il12rb2* expression was analyzed by RT-PCR over the course of the infection. *Il12rb2* mRNA slowly but steadily increased in Treg cells over the first 10 days of LCMV infection, mirroring the acquisition of T-bet (*Tbx21*) in these cells (Fig. 1A) and confirming the expression of *Il12rb2* by Treg cells. To determine whether *Il12rb2* expression increased in the overall Treg cell population or specifically in the type 1 Treg cells marked by T-bet and CXCR3 (*6*), we sorted CXCR3^+^ and CXCR3^−^ Treg cells over the same time period and again analyzed *Il12rb2* and *Tbx21* expression. As expected, *Tbx21* expression was restricted to the CXCR3^+^ Treg cells subset. Similarly, *Il12rb2* was upregulated in CXCR3^+^ but not CXCR3^−^ Treg cells over the course of the infection (Fig. 1B), suggesting that both IL-12R and T-bet are specifically expressed upon type 1 Treg cell differentiation.

**Fig. 1.**
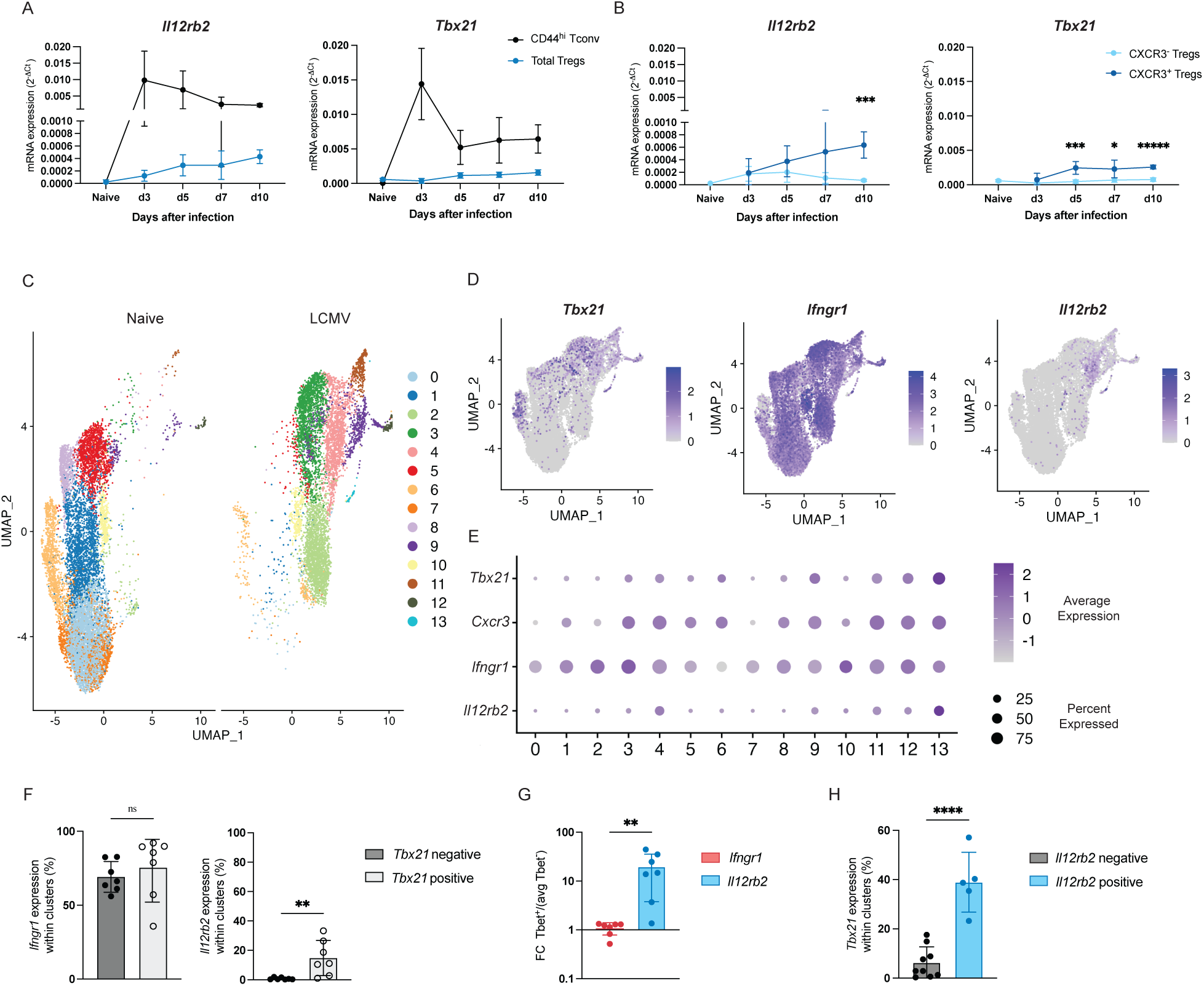
*Il12rb2* is upregulated in T-bet^+^ Treg cells during viral infection. (**A, B**) Foxp3^GFP-KI^ mice were left naïve or infected with LCMV WE 200 f.f.u. Treg cells or CD44^hi^ Tconv were sorted at days 0, 3, 5, 7 and 10 p.i. and quantitative PCR for *Il12rb2* and *Tbx21* was performed. For (B) Treg cells were sorted separately according to CXCR3 expression. (**C-H**) A scRNA-seq dataset containing Treg cells from naive or LCMV-infected mice was reanalyzed. (C) UMAP displaying the Treg clusters separated by condition (infection). Feature plots (D) and dotplot by cluster (E) indicating *Tbx21*, *Cxcr3*, *Ifngr1* and *Il12rb2* expression. (F, G) Treg clusters were divided by *Tbx21* expression. Percentage of cells expressing *Ifngr1* and *Il12rb2* within cluster (F) or fold-change expression between T-bet^+^ and T-bet^−^ clusters (G) are shown. (H) Treg clusters were separated by *Il12rb2* expression and percentage of *Tbx21* expression within clusters is represented. Data in (A, B) are representative from two to three independent experiments per timepoint with n=5-10 mice per group. Corrected multiple t tests (B) and unpaired t tests (F-H) were used. Means with SD are shown. * p < 0.05, ** p < 0.01, *** p < 0.001, **** p < 0.0001, ***** p < 0.00001.

To confirm these results at the single-cell level, we analyzed our single-cell RNA sequencing (scRNA-seq) dataset of sorted Treg cells from naive or LCMV-infected animals(*17*). Dimensionality reduction using Uniform Manifold Approximation and Projection (UMAP) identified 14 clusters, with a clear separation between clusters containing cells retrieved from infected and naive animals (Fig. 1C). In line with the results obtained in the sorted Treg cells populations, we found *Tbx21* and *Cxcr3* to be predominantly expressed in clusters appearing after infection (clusters 3, 4, 9, 11, 12, 13) as well as one naive cluster (cluster 6). Along with other cytokine receptors, *Il12rb2* expression is sparse and difficult to detect in scRNA-seq data.

However, we could consistently detect *Il12rb2* in Treg cells from infected, but not naive, mice (clusters 4, 9, 11, 12 and 13), with concomitant expression of *Tbx21* (Fig. 1D, E). Interestingly, although IFN-γ has been shown to promote type 1 Treg cell differentiation, *Ifngr1* was highly expressed in almost all clusters (Fig. 1D, E). Separating the clusters by *Tbx21* expression revealed that, while *Ifngr1* was not differentially expressed between the two groups, the proportion of cells expressing *Il12rb2* was significantly higher in *Tbx21*-positive than in *Tbx21*-negative clusters (Fig. 1F, G). Moreover, *Tbx21* expression was greatly found in high levels in clusters expressing *Il12rb2* (Fig. 1H). Overall, these data confirm that Treg cells express *Il12rb2* after LCMV infection, that its expression is upregulated specifically in T-bet^+^ Treg cells, and that *Il12rb2* is more closely tied to *Tbx21* expression than *Ifngr1*.

### IL-12Rβ2 KO Treg cells show impaired type 1 specialization after infection

To study the functional role of IL-12Rβ2 in Treg cells, we crossed *Foxp3^cre-YFP^* with *Il12rb2^fl/fl^* mice to generate *Foxp3^cre-YFP^xIl12rb2^fl/fl^* mice, which lack IL-12Rβ2 specifically on Treg cells (*Il12rb2* cKO) and *Cre* activity is reported via YFP expression. We infected *Il12rb2* cKO animals and littermate controls with LCMV and analyzed the phenotype of Treg cells by flow cytometry (fig. S1A). IL-12Rβ2 KO Treg cells showed lower expression of T-bet and CXCR3 in comparison to Treg cells from control animals (Fig. 2A, B). As Foxp3 is encoded by the X chromosome, we used heterogenic *Foxp3^cre-YFP/wt^Il12rb2^fl/fl^* females to study the effect of *Il12rb2* depletion in Treg cells within the same host. We again observed a significant decrease in T-bet and CXCR3 expression in the IL-12Rβ2 KO Treg cells (Fig. 2C, D), confirming the importance of IL-12 signaling for type 1 Treg cell specialization.

**Fig. 2.**
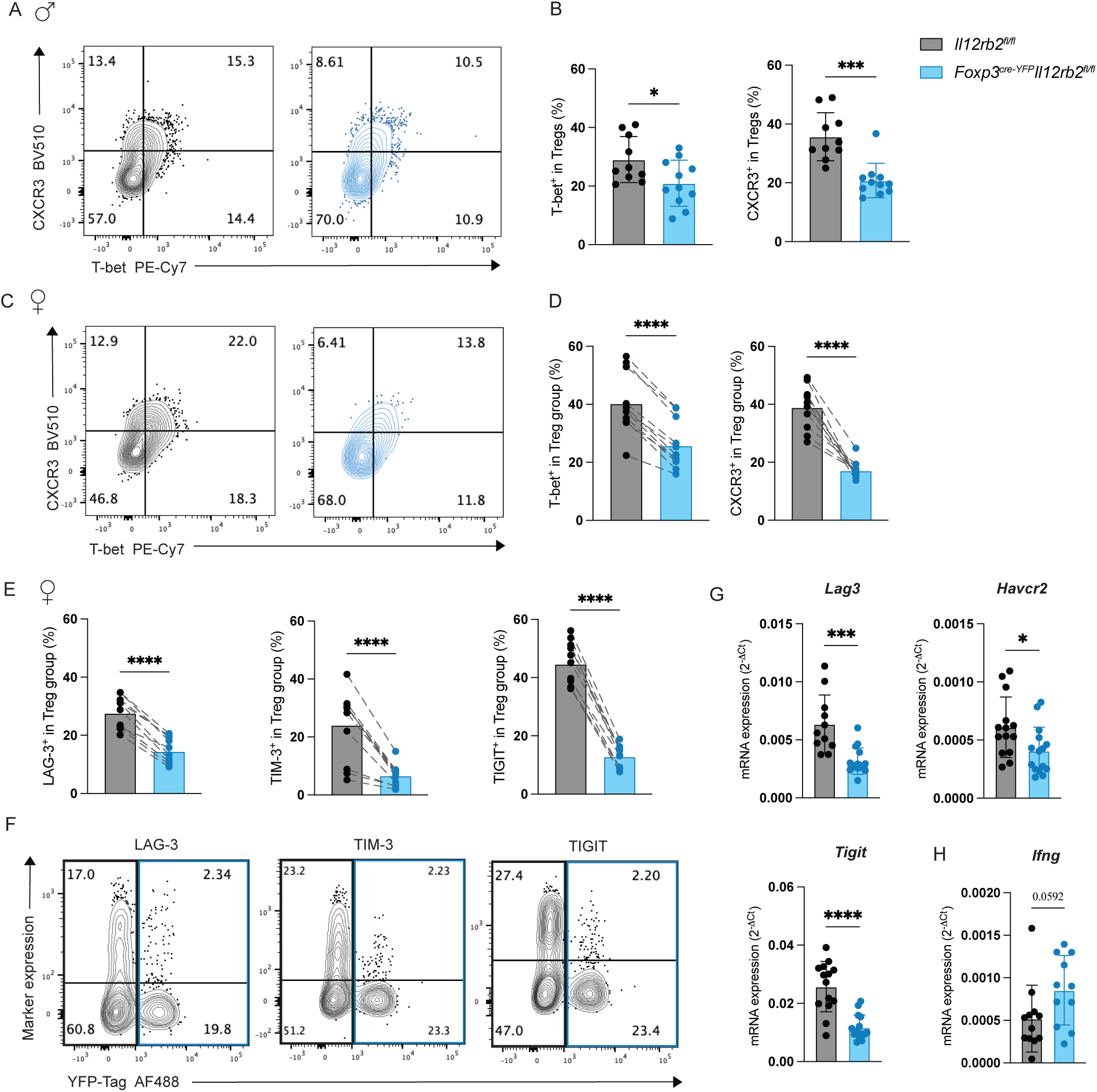
IL-12Rβ2 KO Treg cells display an impaired type 1 specialization. *Foxp3^cre-YFP^Il12rb2^fl/fl^* animals or *Foxp3^wt^Il12rb2^fl/fl^* littermate controls (**A, B**) or heterogenic females *Foxp3^cre-YFP/wt^Il12rb2^fl/fl^* (**C-F**) were infected with LCMV WE 200 f.f.u. At day 10 p.i., Treg cells from spleen were isolated and stained for T-bet, CXCR3 and co-inhibitory receptor expression. (A, C) Representative FACS plots and (B, D) quantification graphs displaying T-bet and CXCR3 expression. (E, F) LAG-3, TIM-3, and TIGIT expression. (**G, H**) *Foxp3^GFP-KI^*(WT) or *Foxp3^cre-YFP^Il12rb2^fl/fl^* mice were infected with LCMV WE 200 f.f.u. At day 10 p.i., Treg cells from spleen were isolated and FACS-sorted, the mRNA isolated and marker expression quantified by RT-qPCR. Data in (A-F) are representative from two independent experiments for each sex with n=10-11 (A, B) and n=11 (C-F) mice per group. Data in (G-H) are pooled from 5 independent sorts with n=11-15 mice per group. Unpaired t test, or paired t test (for D, E), were used. Means with SD are shown. * p < 0.05, *** p < 0.001, **** p < 0.0001.

Our group has previously shown that the co-inhibitory receptors LAG-3, TIM-3 and CD85k are specifically upregulated in Treg cells during Th1 responses(*15*). Furthermore, the coinhibitory receptor TIGIT marks activated Treg cells and its expression is also induced upon LCMV infection(*15*, *18*). We thus used these receptors as phenotypic markers for type 1 Treg cells and found IL-12Rβ2 KO Treg cells to express reduced levels of LAG-3, TIM-3 and TIGIT (but not CD85k) both at the protein and at the mRNA level (Figures 2E-G, fig. S1B), suggesting a defect in their type 1 specialization and function.

We also checked the production of IFN-γ by Treg cells, which remained similar in both groups (Fig. 1H). Thus, contrary to previous reports(*14*), IL-12R signaling does not seem to induce the production of IFN-γ in Treg cells in this context. Taken together, these results suggest that IL-12R signaling in Treg cells contributes in type 1 Treg cells polarization and the acquisition of the co-inhibitory receptors LAG-3, TIM-3 and TIGIT.

### IL-12Rβ2 expression in Treg cells is primarily induced by T-bet downstream the IFN-γ signaling cascade

While IL-12Rβ2 KO Treg cells showed reduced T-bet protein expression, *Tbx21* mRNA levels in these cells were not altered in our experimental setting (Fig. 3A). Hence, we wanted to investigate whether IL-12 could induce *Tbx21* mRNA *in vitro* or whether it rather acts after the mRNA has been transcribed. We cultured FACS-sorted naive CXCR3^−^ Treg cells with different cytokine combinations *in vitro* for 72h and tested for *Tbx21* expression. IL-12, either alone or in combination with IFN-γ, emerged as the most potent inducer of *Tbx21* expression (Fig. 3B).

**Fig. 3.**
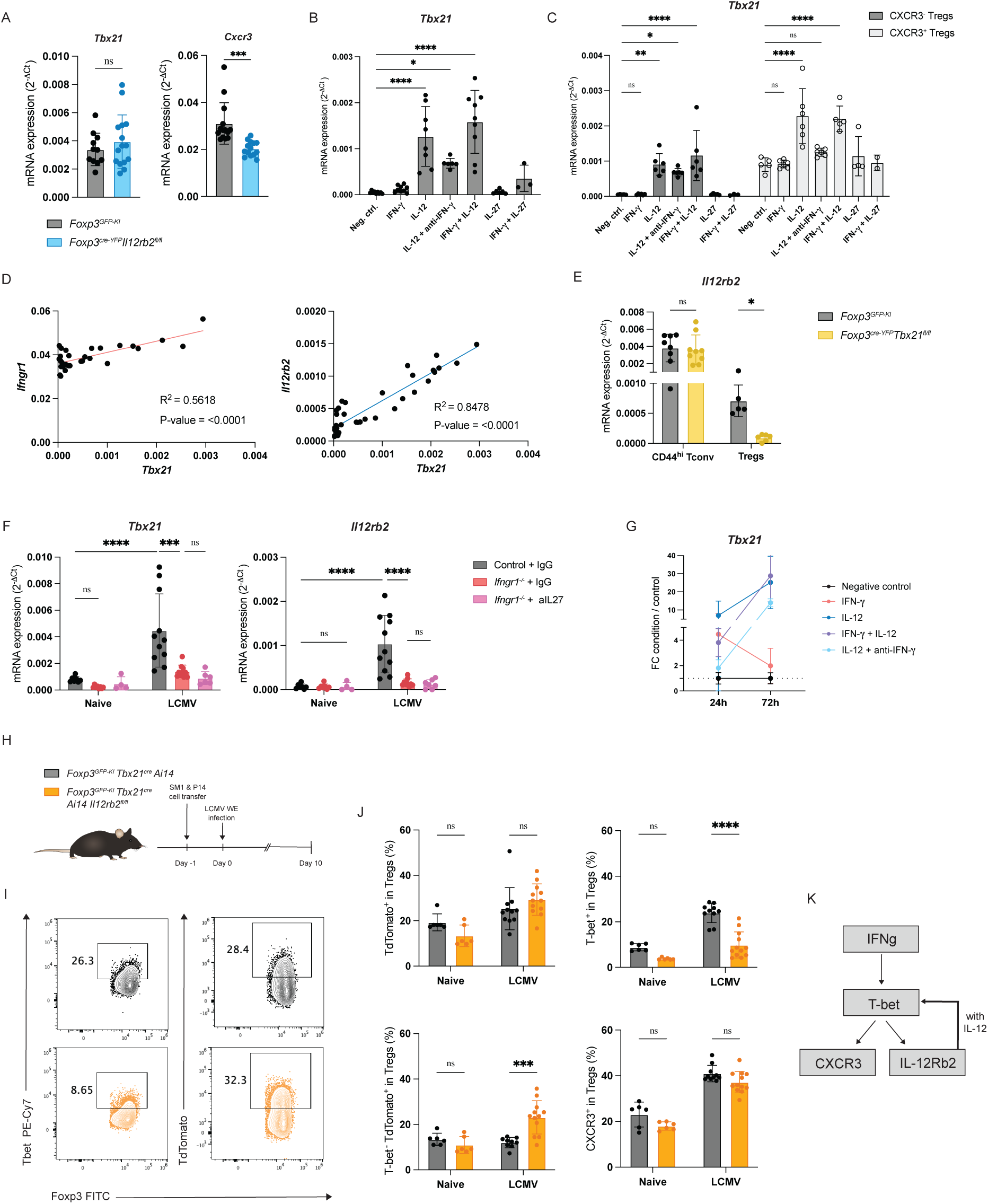
IL-12 signaling acts after IFNγ-driven T-bet activation and reinforces T-bet expression. (**A**) *Tbx21* and *Cxcr3* mRNA expression in total Treg population of *Foxp3^GFP-KI^* (WT) or *Foxp3^cre-YFP^Il12rb2^fl/fl^*mice at day 10 after LCMV WE 200 f.f.u. infection. (**B-D**) CXCR3^−^ Treg cells, or also CXCR3^+^ Treg cells (C), from naive mice were FACS-sorted and incubated with different cytokines for 72h *in vitro* in the presence of IL-2 and anti-CD3 dynabeads. Expression of *Tbx21* mRNA was quantified by RT-qPCR. (D) Correlation plots showing the expression of *Tbx21* vs. *Ifgnr1* or *Il12rb2* in CXCR3^−^ Treg cells after 72h of culture with different cytokines. (**E**) Expression of *Il12rb2* mRNA in *Foxp3^GFP-KI^* or *Foxp3^cre-YFP^Tbx21^fl/fl^*mice at day 10 after infection with LCMV WE 200 f.f.u. (**F**) *Foxp3^GFP-KI^*and *Ifngr1^−/−^* mice were infected with LCMV WE 200 f.f.u. In 2 out of the 4 repeats, a group of *Ifngr1^−/−^* was treated with anti-IL27p28 blocking antibody at days –1, 3 and 7 p.i. In those experiments, the WT and *Ifngr1^−/−^* control groups were treated with IgG2 isotype antibodies. mRNA quantification of *Tbx21* and *Il12rb2* in Treg cells is shown. (**G**) Naïve CXCR3^−^ Treg cells were incubated with different cytokines for 24h or 72h *in vitro* in the presence of IL-2 and anti-CD3 dynabeads. Plot indicating the fold change of *Tbx21* mRNA (dCT cytokine condition divided by dCT of the negative control) expression over time is shown. (**H-J**) *Foxp3^GFP-KI^Tbx21^cre^Ai14* mice were crossed with *Il12rb2^fl/fl^*mice to generate *Foxp3^GFP-KI^Tbx21^cre^Ai14 Il12rb2^fl/fl^*mice. Virus-specific SM1 and P14 cells were transferred a day before infection with LCMV WE 200 f.f.u. (H) Experimental setup. (I, J) T-bet, CXCR3 and TdTomato expression in Treg cells at day 10 p.i. (**K**) Representative model: IFN-γ induces a first peak of T-bet, which drives the expression of CXCR3 and IL12rb2. IL-12 signaling generates a feedforward loop which reinforces T-bet expression after 48h. Data in (A) are pooled from 5 independent sorts with n=11-15 mice per group. Data in (B) are pooled from 3 independent experiments with n=3-9 technical replicates per group. Data in (C) are pooled from 2 independent experiments, different from (B), with n=2-6 technical replicates per group. Data in (D) are analyzed from pooling the CXCR3^−^ samples from (B, C), with n=36. Data in (E) are pooled from 2-3 independent experiments with n=5-9 mice per group. For (F), 4 independent experiments with a total of n=4-11 mice per group. Data in (G) are calculated from 2-5 independent experiments with n=5-15 technical replicates per group. For (H-J), 3 independent experiments with a total of n=6-12 mice per group were used. Unpaired t test (A), one-way ANOVA (B, C), simple linear regression (D) or two-way ANOVA followed by Dunnet’s (c) or Tukey’s (E, F, J) multiple comparisons tests were used. Means with SD are shown. * p < 0.05, ** p < 0.01, *** p < 0.001, **** p < 0.0001.

Previous studies have reported IFN-γ as the main T-bet driver in Treg cells(*14*). However, while we observed a potent induction of *Tbx21* with IL-12, IFN-γ alone failed to elicit an increase. This was not due to lack of IFN-γ sensing, as all Treg cells expressed *Ifngr1* (Fig. 1D, fig. S2A). To test whether this discrepancy could be attributed to differences in the starting Treg cell population, we sorted CXCR3^−^ and CXCR3^+^ Treg cells separately and repeated the assay under the same conditions. While CXCR3^+^ Treg cells displayed higher *Tbx21* baseline expression, both subsets responded similarly to the different cytokine treatments (Fig. 3C). Notably, none of the cytokine stimulations led to noticeable changes in Treg cells proliferation or cell death (fig. S2B). Together, these data indicates that IFN-γ does not induce *Tbx21* mRNA after 72h in culture, whereas IL-12 potently does so.

We next examined the expression of IFN-γR1 and IL-12Rβ2 in our samples. As shown in Fig. 1, *Il12rb2* expression is closely associated with *Tbx21* expression. This correlation was again evident in our *in vitro* cultures, where *Il12rb2* and *Tbx21* expression were clearly correlated (Fig. 3D), suggesting that these two genes are tightly linked. To investigate whether there is a direct link between these receptors and T-bet, we used *Foxp3^cre-YFP^xTbx21^fl/fl^* mice. Upon LCMV infection, T-bet KO Treg cells failed to upregulate *Il12rb2*, identifying T-bet as a critical factor in the induction of IL-12Rβ2 (Fig. 3E), potentially as part of a feedforward loop. Of note, IL-12 was still able to induce *Il12rb2* expression in T-bet KO Treg cells *in vitro*, suggesting that while *Il12rb2* may be downstream of the T-bet *in vivo*, alternative T-bet-independent pathways can also induce its expression (fig. S2C).

Because IFN-γ was reported to induce T-bet and *Ifngr1* is ubiquitously expressed in Treg cells (Fig. 1), we took advantage of *Ifngr1* full KO mice and treated them during the course of the infection with IL-27 blocking antibody or isotype control to study the effect of these cytokines in type 1 Treg cell specialization and IL-12Rβ2 expression. In line with previous findings, *Ifngr1* KO Treg cells showed significantly reduced *Tbx21* after infection(*6*) but also failed to upregulate *Il12rb2* (Fig. 3F). IL-27 could not compensate for the loss of IFN-γ sensing in inducing T-bet (Fig. 3F), although it can be relevant in inducing T-bet in mucosal sites(*13*).

### IL-12 sensing allows for long-lasting T-bet expression in Treg cells

The discrepancy between the ability of IFN-γ to induce T-bet *in vivo* and *in vitro* may reflect temporal changes in its functional role. To explore this, we assessed the kinetics of T-bet expression in response to IFN-γ and IL-12 *in vitro* in sorted WT Treg cells (Fig. 3G, fig. S2D). IFN-γ induced *Tbx21* already 12h post-culture. However, if stimulated with IFN-γ alone, *Tbx21* expression returned to baseline by 48h. In contrast, IL-12 alone (in conditions were IFN-γ was neutralized) had minimal effects during the first 24h of culture, but drove robust *Tbx21* induction by 72h. The combination of both IFN-γ and IL-12 resulted in the highest expression of *Tbx21* mRNA across the entire timecourse. These data suggest that IFN-γ initiates a first, transient wave of *Tbx21* expression in Treg cells that requires IL-12 to be sustained. This first peak of T-bet may be necessary for the upregulation of a fully functional IL-12R *in vivo*, as we observed that IFN-γ together with IL-27 could induce *Il12rb2* in the first 24h (fig. S2E). This would in turn allow for a feedforward loop in which IL-12 reinforces and stabilizes T-bet expression and their type 1 specialization.

To confirm that IL-12 sustains T-bet protein expression after its initial induction, we generated T-bet fate reporter mice, in which *Il12rb2* is deleted upon T-bet induction (*Foxp3^GFP-KI^ Tbx21^cre^ AiRosa^fl/fl^ Il12rb2^fl/fl^*). In these mice, cells constitutively express TdTomato upon activation of the *Tbx21* promoter and simultaneously lose *Il12rb2* expression. We used an acute LCMV infection model to assess T-bet and TdTomato expression in these mice (or in fate reporter controls) and to ensure a functional antiviral response, we transferred LCMV-specific CD4^+^ T cells (Smarta, SM1) and CD8^+^ T (P14) cells one day before infection as the *Il12rb2* depletion is not restricted to Treg cells in these model (Fig. 3H). Strikingly, Treg cells depleted of *Il12rb2* showed a trend of higher frequencies of TdTomato^+^ cells compared to WT controls, indicating robust *Tbx21* induction (Fig. 3I, J). However, we found T-bet (and a trend in CXCR3) protein levels to be decreased in the absence of IL-12Rβ2, indicating that while initial T-bet induction occurs, IL-12R signaling is required to maintain T-bet expression (Fig. 3J). Together, these results support a two-step model in which IFN-γ drives a first peak of T-bet, leading IL-12Rβ2 upregulation, while IL-12 is required beyond 48h to stabilize T-bet expression and sustained type 1 Treg cell specialization (Fig. 3K).

### IL-12Rβ2 KO Treg cells have a reduced capacity to suppress effector T cell expansion and cytokine production

Given that IL-12Rβ2 KO Treg cells show decreased co-inhibitory receptor expression, we hypothesized that their suppressive capacity might be compromised. To test this, we performed *in vitro* suppression assays using activated CD44^hi^ Tconv cells isolated from LCMV-infected mice as responders in co-cultures with either WT or IL-12Rβ2 KO Treg cells (Fig. 4A). We detected a higher proliferation of Tconv cells when co-cultured with IL-12Rβ2 KO Treg cells, and thus, reduced suppressive function in these cells (Fig. 4B, C). Furthermore, sorted CD44^hi^ Tconv isolated from LCMV-infected *Foxp3^cre-YFP^xIl12rb2^fl/fl^* mice also showed higher expression of *Tbx21* and other activation markers at the mRNA level compared to control mice (fig. S3A).

**Fig. 4.**
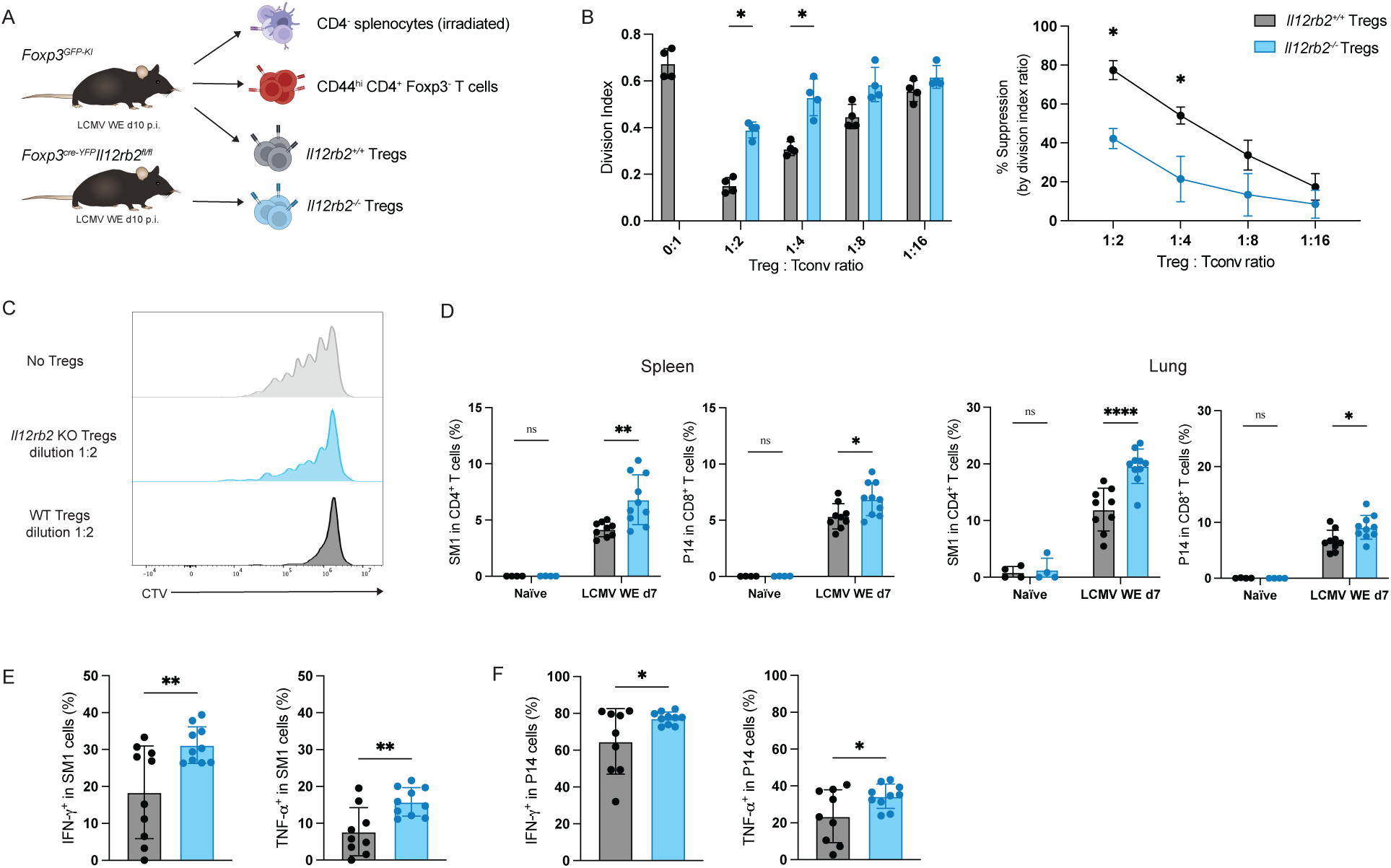
IL-12Rβ2 KO Treg cells are less suppressive. (**A-C**) Suppression assay performed with WT or IL-12Rβ2 KO Treg cells. CD44^hi^Foxp3^−^CD4^+^ T conventional cells (Tconv) and Foxp3^+^CD4^+^Treg cells were FACS-sorted at day 10 after infection with LCMV WE 200 f.f.u. Tconv isolated from WT mice and labelled with CTV were seeded with different ratios of WT or IL-12Rβ2 KO Treg cells, in the presence of irradiated CD4^−^ splenocytes (as APCs) and anti-CD3. 72h later, cell proliferation was measured using a spectral cytometer. (A) Experimental setup. (B) Tconv proliferation and percentage of suppression measured by Cell-Trace Violet (CTV) dilution. (C) Representative histograms of CTV expression in Tconv in the 1:2 condition. (**D-F**) 2.500 SM1 and 5.000 P14 cells were transferred to *Foxp3^cre-YFP^Il12rb2^fl/fl^* animals or *Foxp3^wt^Il12rb2^fl/fl^* littermate controls one day prior infection with LCMV WE 200 f.f.u. (D) Percentage of SM1 cells among CD4^+^ T cells and P14 cells among CD8^+^ T cells in spleen or lung at day 7 p.i. (E) Production of IFN-γ and TNF-α by SM1 or (F) P14 cells at day 7 p.i. Data in (B,C) are representative from 1 out of 2 independent experiments with n=4 technical replicates per condition. Data in (D-F) are pooled from 2 independent experiments with n=4-10 mice per group. Corrected multiple t tests (B), two-way ANOVA followed by Tukey’s multiple comparisons tests (D) and unpaired t tests (E, F) were used. Means with SD are shown. * p < 0.05, ** p < 0.01.

To assess their suppressive capacity *in vivo*, we transferred LCMV-specific CD4^+^ T cells (Smarta, SM1) and CD8^+^ T cells (P14) into WT or IL-12Rβ2 cKO animals, followed by LCMV infection. Seven days post infection, we determined the expansion of the transferred cells in spleen and lung. In line with our *in vitro* findings, SM1 and P14 cells showed significantly greater expansion in IL-12Rβ2 cKO mice, confirming that IL-12Rβ2 KO Treg cells are less effective in suppressing T cell proliferation *in vivo* (Fig. 4D). In addition to the increased proliferation, both effector populations also showed an enhanced production of IFN-γ and TNF-α (Fig. 4E). In conclusion, these results demonstrate that IL-12R signaling in Treg cells contributes in efficiently controlling CD4^+^ and CD8^+^ T cell proliferation and cytokine production.

### Lack of IL-12Rβ2 in Treg cells alters CD8^+^ T cell exhaustion

Functional Treg specialization also occurs during chronic infections(*19*). Seeing how IL-12Rβ2 KO Treg cells are less suppressive, we wanted to explore whether the impact of IL-12 on Treg specialization also translates to changes during chronic LCMV infection, specifically on the CD8^+^ T cell compartment. As expected upon a chronic infection, both *Foxp3^cre-YFP^Il12rb2^fl/fl^* and control mice showed an increase in Treg cells after infection with LCMV Clone 13 (Fig. 5A). However, similar to our observations during acute infection, IL-12Rβ2 KO Treg cells showed a markedly reduced T-bet and CXCR3 expression 27 days post-infection (Fig. 5B). Additional Treg specialization markers such as LAG-3, TIM-3 and TIGIT (but not CD85k) were also diminished in Treg cells from IL-12Rβ2 cKO animals (Fig. 5C). Thus, Treg specialization is maintained throughout the course of a chronic infection and IL-12R signaling is required to support this program across both acute and chronic settings.

**Fig. 5.**
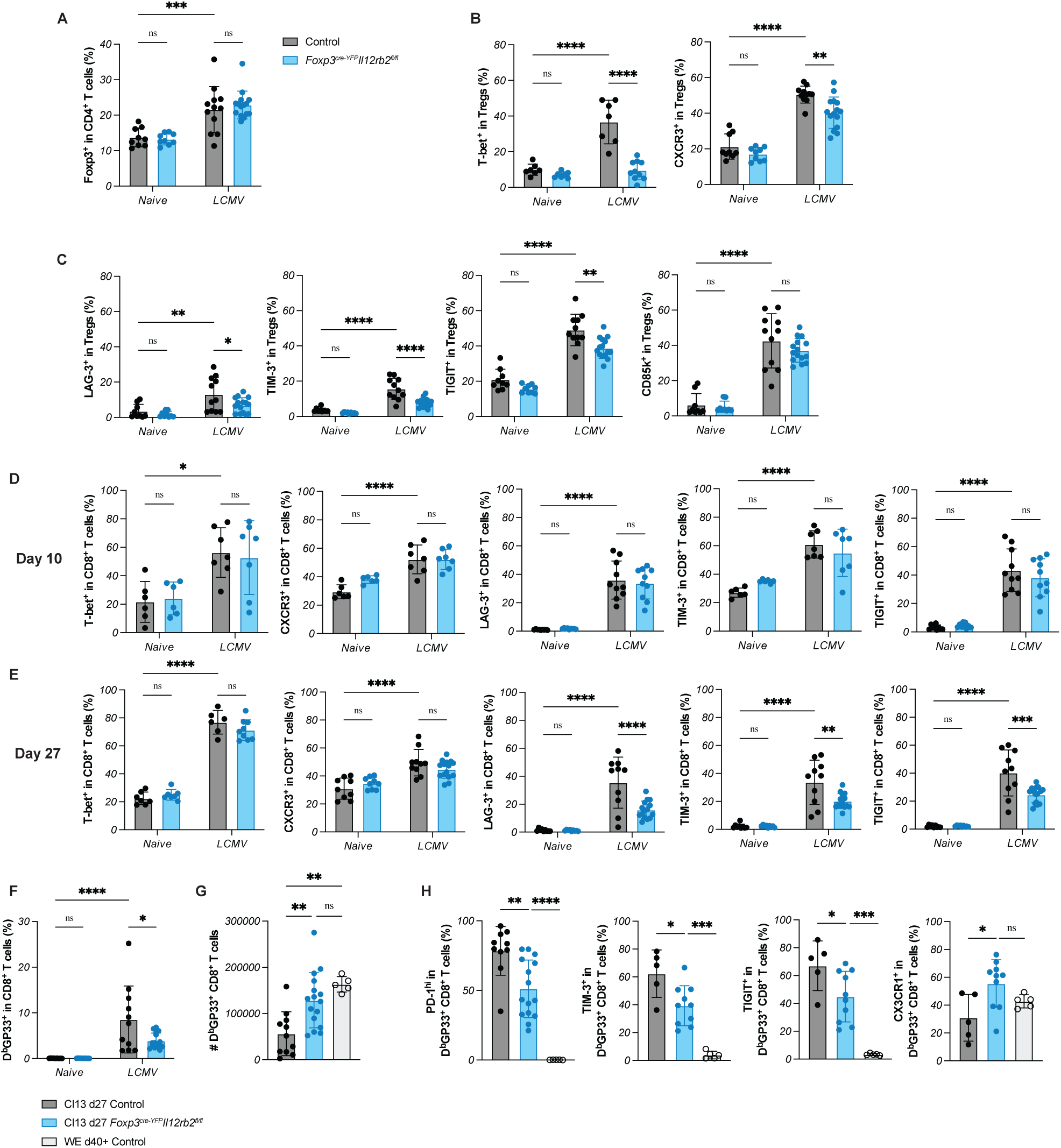
CD8^+^ T cell exhaustion is altered by IL-12Rβ2 KO Treg cells. (**A-C, E-I**) *Foxp3^cre-YFP^Il12rb2^fl/fl^*mice or littermate controls were infected with LCMV Cl13 2Mio f.f.u. and splenocytes isolated at day 27 p.i. (A) Percentage of Treg cells among CD4^+^ T cells. (B) Treg expression of T-bet and CXCR3 or (c) the co-inhibitory receptors LAG-3, TIM-3, TIGIT and CD85k. (**D**) 2.500 SM1 and 5.000 P14 cells were transferred a day prior to infection with LCMV Cl13 2×10^6^ f.f.u. Expression of markers in host CD8^+^ T cells at day 10 p.i. is shown. (E) Expression of markers in host CD8^+^ T cells at day 27 p.i. (F) Percentage and (G) absolute numbers of DbGP33^+^ cells in spleen. (H) Expression of PD-1^hi^, TIM-3, TIGIT and CX3CR1 in DbGP33^+^ CD8^+^ T cells. Data are pooled from 2-3 independent experiments with n=5-16 animals per group. Two-way ANOVA (A-F) or one-way ANOVA (G, H) followed by Tukey’s multiple comparisons tests were used. Means with SD are shown. * p < 0.05, ** p < 0.01, *** p < 0.001, **** p < 0.0001.

In CD8^+^ T cells, T-bet and CXCR3 expression were increased upon infection in WT and IL-12Rβ2 cKO mice. However, on day 27 post-infection, the expression of co-inhibitory receptors on these cells was reduced in IL-12Rβ2 cKO animals (Fig. 5E). Notably, these differences were not observed at earlier timepoints (day 10, Fig. 5D), suggesting that the altered phenotype is not due to a decreased activation, but rather an atypical T cell exhaustion process. To confirm that these results were not merely an artifact of reduced activation, we further analyzed DbGP33^+^ LCMV-specific CD8^+^ T cells. While the frequency of DbGP33^+^ CD8^+^ T cells after LCMV infection was slightly reduced in IL-12Rβ2 cKO animals (Fig. 5F), their total number was greater than in WT animals and reached levels comparable to those observed in memory mice that had cleared acute LCMV infection (Fig. 5G). This increase in numbers was a consequence of increased total cell counts in IL-12Rβ2 cKO mice, suggesting that these mice are better protected from lymphopenia. In fact, other groups also observed an increase in splenocyte count during chronic LCMV infection after Treg cells modulation(*20*). Similarly, while percentages of IFN-γ-expressing DbGP33^+^ CD8^+^ T cells remained similar between groups, IL-12Rβ2 cKO mice harbored significantly higher total numbers of these cells (fig. S4A), which translated into a decrease in viral titers in spleens of IL-12Rβ2 cKO mice (Fig. S4B).

The proportion of PD-1^hi^ DbGP33^+^ cells, as well as TIM-3 and TIGIT expression, were reduced in mice lacking IL-12Rβ2 in Treg cells, indicating a less exhausted CD8^+^ T cell phenotype (Fig. 5H). Furthermore, “effector-like” exhausted cells, marked by CX3CR1 expression(*21*), were increased in IL-12Rβ2 cKO animals. In summary, lack of IL-12 sensing in Treg cells increases the total pool of virus-specific CD8^+^ T cells and changes their exhaustion state towards a less exhausted phenotype.

### IL-12 contributes to type 1 specialization in human Treg cells

Finally, we wanted to investigate whether these findings could also be translated to human Treg cells. We isolated CD4^+^CD127^low^CD25^+^ CXCR3^−^ Treg cells from peripheral blood mononuclear cells (PBMCs) of healthy donors and cultured them with cytokines for 120h (fig. S5A). Human Treg cells showed mild induction of T-bet and CXCR3 when incubated with IFN-γ (Fig. 6A). However, in line with our findings in mouse models, IL-12 was a much stronger inducer of T-bet and CXCR3 in human Treg cells. The combination of IL-12 with IFN-γ did not further enhance *Tbx21* expression beyond IL-12 alone, indicating that in this setting IFN-γ plays a minimal role. Notably, human Treg cells are capable of expressing *IL12RB2* at steady-state conditions(*22*), which may explain their diminished dependence on IFN-γ. Moreover, other groups also reported the inability of IFN-γ in inducing IL-12Rβ2 in human Th1 cells(*23*).

**Fig. 6.**
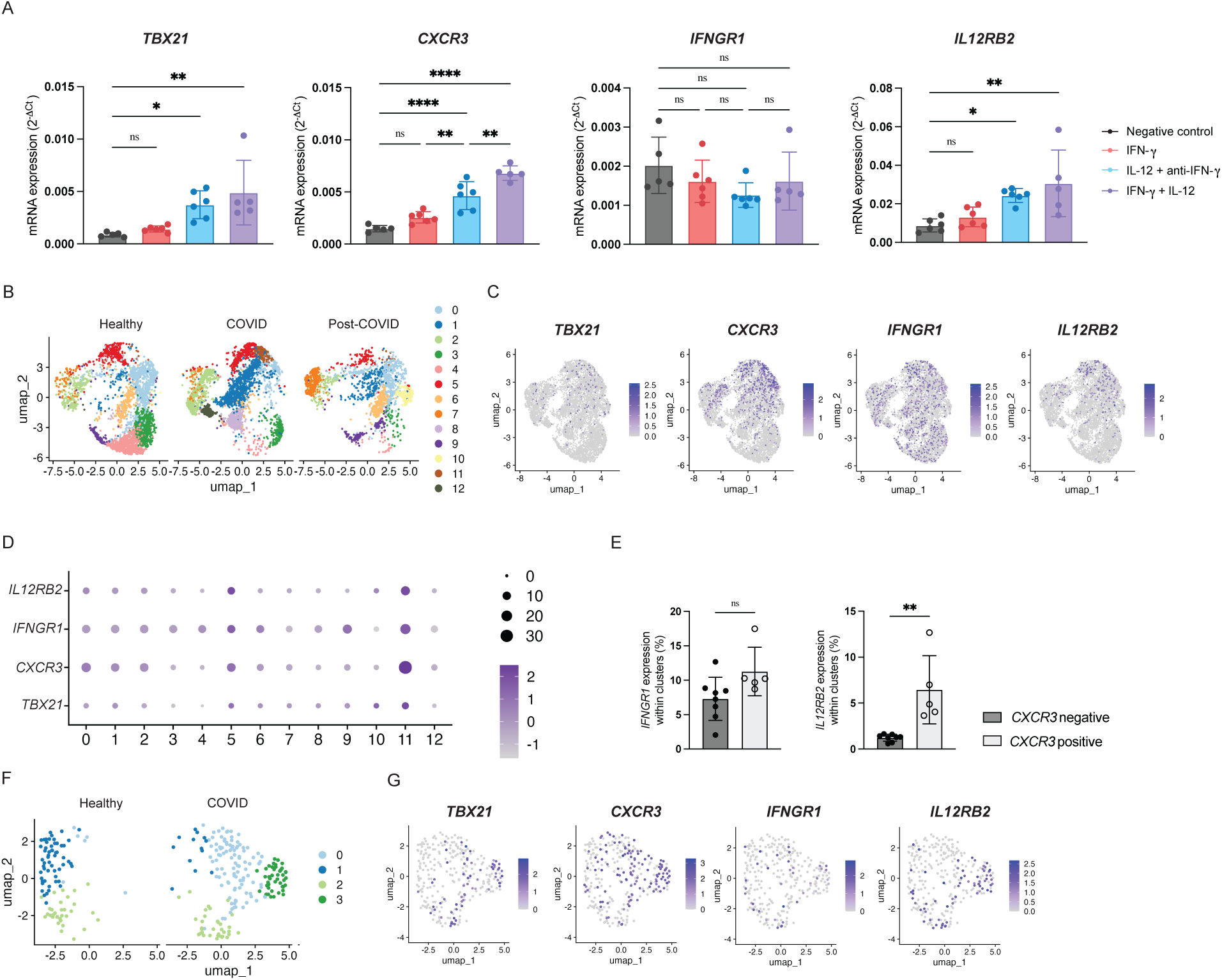
IL-12 contributes to type 1 specialization in human Treg cells. (**A**) CXCR3^−^ Treg cells isolated from PBMCs of healthy donors were incubated with IL-2 and anti-CD3 and indicated experimental conditions. Expression of *TBX21*, *CXCR3, IFNGR1 and IL12RB2* were measured at day 5. (**B-G**) scRNA-seq data from COVID patients or healthy donors was reanalyzed. (B-E) Treg cells from PBMCs were subsetted for downstream analysis. (B) UMAP of Treg clusters by disease stage, feature plots (C) and dotplot by cluster (D) indicating *TBX21*, *CXCR3*, *IFNGR1* and *IL12RB2* expression. (E) Expression of *IFNGR1* and *IL12RB2* in *CXCR3*-negative vs. *CXCR3*-positive clusters. (F) UMAP of Treg clusters by disease stage and (G) feature plots for *TBX21*, *CXCR3*, *IFNGR1* and *IL12RB2* expression in Treg cells isolated from nasopharyngeal aspirates. For (A), data are pooled from 3 independent experiments with 3 different donors with n=5-6 technical replicates per group. One-way ANOVA (A) followed by Tukey’s multiple comparisons tests or unpaired t tests (E) were used. Means with SD are shown. * p < 0.05, ** p < 0.01, **** p < 0.0001.

To explore Treg specialization in the context of an ongoing immune response, we took advantage of a publicly available scRNA-seq dataset from COVID-19 patients(*24*). This dataset includes PBMCs and lung aspirates of patients during or after Sars-CoV-2 infection, as well as healthy controls. We identified 13 Treg cells clusters in the PBMCs (Fig. 6B), with type 1 markers primarily expressed in clusters from cells retrieved during COVID-19 (Fig. 6C). Clusters 5 and 11 were the main clusters expressing *IL12RB2* and these also showed the highest expression of *TBX21* and *CXCR3* (Fig. 6D). In line with what we observed in our murine scRNA-seq data, *IFNGR1* was broadly expressed across clusters. Due to relatively low *TBX21* expression in this dataset, we used CXCR3 as a proxy for T-bet^+^ Treg cells and stratified clusters based on CXCR3 expression. While *IFNGR1* expression did not differ between *CXCR3-*positive and negative clusters, *IL12RB2* expression was significatnly higher in *CXCR3-*expressing clusters (Fig. 6E). These results indicate that *IL12RB2* is co-expressed with T-bet in human Treg cells during an ongoing immune response. In lung aspirates, 4 Treg cells clusters were identified, which also clearly showed coordinated expression of *TBX21*, *CXCR3* and *IL12RB2* among clusters (Fig. 6G). Overall, human Treg cells can undergo type 1 specialization in response to infection and IL-12 plays a central role in driving this process.

## Discussion

In this study, we identify IL-12 as a critical driver of type 1 Treg cell specialization and suppressive function. While previous work described IFN-γ as the main inducer of Treg specialization, our data suggests that the key role of IFN-γ is to induce IL-12R expression in Treg cells, thereby enabling IL-12-dependent maintenance of the specialized Treg cell phenotype.

Using LCMV infection as a model for type 1 immune responses, we observed a gradual induction of *Il12rb2* in Treg cells after infection, paralleling *Tbx21* expression. Unlike *Ifngr1*, which was broadly expressed in Treg cells, *Il12rb2* was restricted to T-bet^+^ clusters that emerged after LCMV infection. Sorting Treg cells by CXCR3 expression (as a surrogate marker for T-bet)(*25*) confirmed that *Il12rb2* is specific to type 1 Treg cells. Our data suggests that the critical role of IFN-γ in Treg specialization lays in its ability to induce a transient wave of T-bet, which in turn enables IL-12R expression. Consistent with mechanisms described in Th1 cells, where *Il12rb2* is induced by IFN-γR and/or IL-27R(*26*) signaling(*11*, *27*), we found that during Th1-dominated responses, *Ifngr1* KO Treg cells completely lack *Tbx21* and *Il12rb2* expression. Similarly, T-bet can induce IL-12Rβ2 in Th1 cells(*10*, *28*) and we observed that T-bet KO Treg cells totally lack *Il12rb2* expression *in vivo*. Furthermore, while IFN-γ was able to induce a first peak of T-bet in naive Treg cells *in vitro*, this could not be sustained unless IL-12 was added. Thus, our data support a model in which IFN-γ primes Treg cells for type 1 specialization, while IL-12 consolidates and maintains this phenotype both *in vitro* and *in vivo*.

Our experiments using fate reporter mice further emphasize the importance of IL-12R signaling after T-bet engagement. Deletion of *Il12rb2* after T-bet induction impaired maintenance of T-bet expression in Treg cells, despite a slight increase in reporter-positive Treg cells in those animals. This compensatory response emphasizes the importance of Treg specialization in this context and the critical role of IL-12 in this process. Functionally, IL-12Rβ2 KO Treg cells loose suppressive capacity both *in vitro* and *in vivo*, which goes along with reduced expression of the co-inhibitory receptors LAG-3, TIM-3 and TIGIT. Furthermore, these Treg cells have a decreased CXCR3 expression, which could be required for migration and interaction with their target cells. While the differences in CXCR3 expression did not contribute to Treg cells suppression *in vitro*, it may compromise recruitment and positioning of type 1 Treg cells *in vivo*, especially in localized immune challenges.

Of note, the LCMV model induces a strong type 1 Treg cell specialization and Th1 response that is dominated by type I IFN but only a modest IL-12 response, a fact that restricted the potential of IL-12R signaling in Treg cells. Nevertheless, ablation of these limited IL-12 signal was sufficient to disrupt Treg specialization, phenotype and function.

T-bet^+^ Treg cells not only suppress Th1 responses but also regulate the function of CD8^+^ T cells. In fact, other groups described that type 1 Treg cells regulate the dynamics of CD8^+^ T cells subsets and facilitate the generation of tissue-resident memory T cells(*29–31*). In chronic infections, impaired Treg specialization altered the CD8^+^ T cell compartment to a more “memory-like” phenotype. Virus-specific CD8^+^ T cells from IL-12Rβ2 cKO mice show decreased co-inhibitory receptor expression. These mice also showed a two-fold expansion of gp33-specific CD8^+^ T cells. While slightly lower than the previously reported 10 to 100-fold expansion following a complete loss of Treg cells(*32*), this finding highlights the importance of IL-12-driven Treg cell specialization in limiting the immune response during chronic infections. In line with previous reports that suggest that Treg cells maintain CD8^+^ T cell exhaustion(*32*), the Treg cell-driven changes in CD8^+^ T cell expansion and phenotype translated into slightly reduced viral loads. Whether these effects result from Treg cells shaping the exhausted CD8^+^ T cells directly or indirectly through suppression of CD4^+^ T cell help remains to be determined.

A current debate in the field concerns whether IFN-γ production by Treg cells reflects instability or, alternatively, functional specialization(*33*). In murine models, IL-12 can induce IFN-γ expression in Treg cells(*34*), and IFN-γ production by Treg cells has been suggested as a marker for active, type 1–specialized Treg cells that contribute to the control of Th1 responses(*13*). In contrast, human Treg cells expressing *Il12rb2* can acquire a Th1 effector-like phenotype in autoimmune diseases, a trait that was accompanied with lower suppressive capacities(*35*). Interestingly, we found IL-12Rβ2 KO Treg cells from LCMV-infected mice had even slightly higher *Ifng* transcript levels than WT Treg cells, demonstrating that IL-12 is not the sole driver of IFN-γ expression in Treg cells. Importantly, these IL-12Rβ2 KO Treg cells were less effective in suppression assays, indicating that IFN-γ expression alone does not define functional specialization. Rather, our findings suggest that in murine Treg cells, IL-12R signaling is required to consolidate type 1 specialization and enable efficient suppression of Th1 effector responses, irrespective of IFN-γ expression.

The dependence of Treg cells on IFN-γ and IL-12 could reflect a broader regulatory feedback loop by which Treg cells tune the magnitude of the Th1 response. IFN-γ, but not IL-12, is produced by effector T cells (and NK cells)(*36*). We propose that dendritic cells (DCs) drive both Th1 and type 1 Treg cells polarization(*37*). Indeed, Hall et al. reported that DCs from infected mice produced IL-12 and IL-27, inducing T-bet expression in Treg cells during *Toxoplasma gondii* infection(*13*). Given that DC abundance influences Treg cells numbers(*38*), DCs may be key players in determining the magnitude and phenotype of Treg cells responses.

Functional Treg specialization might allow for targeting of specific Treg cells subsets without impairing general immune homeostasis. Our study shows that IL-12R signaling in Treg cells contributes to T-bet expression and type 1 Treg cell specialization. Impairment in IL-12R signaling renders Treg cells less suppressive and alters the ongoing immune response. As human Treg cells can express *Il12rb2* at steady-state(*22*), modulating Treg cell phenotype with targeted IL-12 therapies could be exploited in type 1-dominated diseases.

## Materials and Methods

### Study Design

This study was designed to investigate the role of IL-12R signaling in type 1 Treg cell specialization across multiple molecular levels. To this end, two conditional KO mouse lines were generated, carrying a deletion of the *Il12rb2* gene under the control of Foxp3 or T-bet expression. Treg cell phenotypes were analyzed at the transcriptional and protein levels. I addition, we assessed whether alterations in Treg cell phenotype translated into altered suppressive function *in vitro* and during viral infections *in vivo*. All experiments were performed as 2 to 3 independent repeats, with age– and sex-matched animals across experimental groups.

### Mice, pathogens and infections

All animal experiments were reviewed and approved by the cantonal veterinary office of Zurich (licenses ZH196/2020, ZH219/2020, ZH200/2023, ZH022/2024) and performed according to Swiss legislation. Foxp3-CRE (JAX stock #016959)(*39*), *Il12rb2^fl/fl^* (*40*), *Foxp3*-GFP.KI(*41*), Smarta(*42*) and P14(*43*) TCR transgenic mice were described previously. Tbet-Tref mice were obtained by crossing Ai14 (JAX stock #007914), T-bet-CRE (JAX Stock #024507) and *Foxp3*-GFP.KI(*41*). Tbet-Tref was further crossed with *Il12rb2^fl/fl^* (*40*). Mice were bred and housed in SPF or OHB facilities of the Laboratory Animal Sciences Center (LASC) Zurich, Switzerland.

Lymphocytic choriomeningitis virus (LCMV) WE was propagated on L929 fibroblasts and LCMV Clone13 (Cl13) on BHK-21 cells, as described previously(*18*). Sex– and age-matched mice, 7-14 weeks of age, were infected i.v. with 200 f.f.u. LCMV WE or 2×10^6^ f.f.u. LCMV Cl13.

### Anti-IL27p28 treatment

To study the effect of combined IFN-γ and IL-27 depletion, in 2 out of the 4 repeats, a group of *Ifngr1^−/−^* was treated with 250µg of anti-IL27p28 blocking antibody at days –1, 3 and 7 p.i. In those experiments, WT and *Ifngr1^−/−^* control groups were treated with IgG2a isotype antibodies. Antibody information is listed in table S1.

### Adoptive cell transfers

Congenitally marked CD45.1^+^ CD4^+^ T cells from Smarta mice and CD45.1^+^ CD8^+^ T cells from P14 mice were purified from splenocytes by negative selection using Mojosort Mouse CD4 or CD8 T cell isolation kits (BioLegend), respectively. Cells were diluted in PBS and 2.5×10^3^ Smarta cells and 5×10^3^ P14 cells were transferred i.v. one day prior to infection. Details for the isolation kits used are listed in table S1.

### Flow cytometry and cell sorting

Splenocytes were collected by mechanical disruption of the spleen by using plunger and a metal mesh. Single cell suspensions were then incubated 3-5min with ACK Buffer (155 mM NH_4_Cl, 1 mM KHCO_3_, 0.1 mM NA_2_EDTA in ddH_2_O, pH 7.2–7.4) for red blood cell lysis. Lung immune cells were isolated using a gentleMACS dissociator (Miltenyi Biotec) and enzymatic digestion with Collagenase I (2.4mg/ml, Roche) and DNase I (0.2 mg/ml, Roche) for 30 min, followed by a 30% Percoll (GE Healthcare) gradient.

For flow cytometry, cell suspensions were resuspended in RPMI medium with 10% FBS, penicillin (50 U/ml, Gibco), streptomycin (50 µg/ml, Gibco) and 2 mM glutamine (Gibco) and, for cytokine analysis, restimulated with the immunodominant LCMV peptides gp33 (1ug/ml, KAVYNFATM) and gp61 (1ug/ul, GLKGPDIYKGVYQFKSVEFD) or αCD3 (1ug/ul, BioXcell) for 4h at 37°C in the presence of Brefeldin A (1ug/ml).

Extracellular staining was performed by diluting antibodies in PBS and incubating the samples for 20min at room temperature (RT). For transcription factor staining, cells were fixed for 40min at RT using the Foxp3 Fixation Buffer (Invitrogen). For assessing cytokine production, a 10 min incubation using BD Cytofix/Cytoperm (BD Biosciences) was used instead. Intracellular staining was performed by diluting the antibodies in Permeabilization buffer (eBiosciences) and incubation for 30min at RT. Gp33-MHC monomers were produced as previously described(*44*) and tetramerized using Streptavidin-APC (BioLegend). Samples were measured using the cytometers BD Fortessa, BD Symphony or Cytek Aurora.

For cell sorting, CD4^+^ T cells were enriched by positive selection using Mojosort Mouse CD4 nanobeads (BioLegend), according to the manufacturer’s instructions. Afterwards, cells were resuspended in supplemented RPMI (10% FBS, 1%PSG) and stained for 30min on ice. Cells were sorted using RPMI medium (2% FBS, 1%PSG) and collected in RPMI 20% FBS for optimal cell viability. Treg cells were sorted by their *GFP* or *YFP* expression. In *Ifngr1^−/−^*, Treg cells were sorted by CD44^mid^ CD25^hi^ expression and *Foxp3^GFP-KI^* animals were used to confirm the sorting strategy. The desired populations were sorted using the 70um or 85um nozzle in BD Aria or BD S6 cell sorters. Details for flow cytometry reagents and antibodies used are listed in table S1.

### Reverse transcription quantitative PCR (RT-qPCR)

The mRNA was isolated using an RNA purification kit (Qiagen) according to manufacturer’s instructions. Reverse transcription was performed using the High Capacity cDNA Kit (Life Technologies). Thermal cycling was performed with a QuantStudio 5 real-time PCR system (ThermoFischer). Desired genes were measured using primer-probe mixtures in FAM-MGB (Applied Biosystems), while normalization was performed by measuring in-sample beta-actin in VIC-MGB (Applied Biosystems). For TIGIT, the following primers and probe were used: forward primer: 5′-CTGATACAGGCTGCCTTCCT-3′, reverse primer: 5′-TGGGTCACTTCAGCTGTGTC-3′, probe: 5′-AGGAGCCACAGCAGGCACGA-3′ (FAM, TAMRA). Details for reagents and probes used are listed in table S1.

### Viral titer quantification

Organs were collected in Minimum Essential Media (MEM, Gibco) supplemented with 2% FBS, penicillin (50 U/ml, Gibco), streptomycin (50 µg/ml, Gibco) and 2 mM glutamine (Gibco) in tubes containing a metal bead. Organs were homogenized using a TissueLyser II (Qiagen). Viral RNA isolation was performed using QIAamp Viral RNA mini Kit (Qiagen), and reverse transcription and RT-qPCR were performed as detailed in the previous section. The following primers and probes were used: forward primer: 5′-ACTGACGAGGTCAACCCGG-3′, reverse primer: 5′-CAAGTACTCACACGGCATGGA-3′, probe: 5′-CTTGCCGACCTCTTCAATGCGCAA-3′ (FAM, TAMRA). Details for reagents and probes used are listed in table S1.

### In vitro Treg cells assays

Treg cells (2.5×10^5^ cells/ml), or Th cells (10^6^ cells/ml) as controls, were cultured *in vitro* with anti-CD3/CD28 stimulation (4×10^4^ beads/well, ThermoFischer), IL-2 and different cytokine combinations (IFNγ, IL-12, IL-27) cultured in Treg cells medium (RPMI medium supplemented with 10% FCS, 50 mM β-mercaptoethanol, 1 mM sodium pyruvate (Gibco), non-essential amino acids (Gibco), 25mM HEPES (Gibco), MEM vitamins (Gibco), penicillin (50 U/ml, Gibco), streptomycin (50 µg/ml, Gibco), and 200 mM glutamine (Gibco)). The final cytokine concentrations were the following: IL-2 (100U/ml), IL-12 (10ng/ml), IL-27 (20ng/ml) and IFN-γ (25ng/ml). In some conditions, anti-IFN-γ (XMG1.2, 10ug/ml, BioLegend) was added. Details for reagents used are listed in table S1.

### In vitro suppression assays

Splenocytes were enriched for CD4^+^ T cells using Mojosort Mouse CD4 nanobeads (BioLegend). The negative fraction was collected, irradiated (36 Gy) and used as APCs. FACS-sorted CD44^hi^ Tconv cells (4×10^4^ cells/well) were cultured in the presence of irradiated splenic APCs (2×10^5^ cells/well) and soluble αCD3 (1ug/1ml, BioXcell) with titrated amounts of Treg cells. Prior seeding, Tconv cells were labelled with CellTrace Violet (ThermoFischer) according to the manufacturer’s protocol. 72h after seeding, cells were harvested and proliferation was assessed by flow cytometry. Details for reagents used are listed in table S1.

### Assays with human cells

Blood was collected from anonymous healthy donors in accordance with Swiss regulations and approved by the Cantonal Ethics Committee of Zurich (BASEC 2024-01233) following written informed consent. In brief, venous blood from healthy participants was collected in EDTA Vacutainer tubes (BD Biosciences) and PBMCs isolated by centrifugation gradient using Ficoll Paque (Cytiva) according to standard protocols. After using CD14 Microbeads (Miltenyi), T cells were purified from the unlabelled fraction using CD4 MicroBeads (Miltenyi) and stained for sorting as outlined for murine T cells. Right before the sort, 7-ADD Viability Staining Solution (BioLegend) was added to distinguish live cells. Tconv cells (CD3^+^CD4^+^CD25^−^CD45RA^+^CXCR3^−^) and Treg cells (CD3^+^CD4^+^CD127^−^CD25^+^CXCR3^−^) were sorted using the 85um nozzle in BD Aria or BD S6 cell sorters. 5×10^4^ Treg cells/well or 2.5×10^5^ Tconv/well were cultured *in vitro* for a total of 5 days in the presence of anti-CD3/CD28 stimulation (8×10^4^ beads/well, Gibco), IL-2 (5ng/ml, R&D Systems) and different cytokine conditions in supplemented RPMI (10% FBS, 50 mM β-mercaptoethanol, 1 mM sodium pyruvate (Gibco), 25mM HEPES (Gibco), 10mM non-essential amino acids (Gibco), penicillin (50 U/ml, Gibco), streptomycin (50 µg/ml, Gibco), and 200 mM glutamine (Gibco)). For Tconv, IL-7 (4ng/ml, R&D Systems) was added instead of IL-2. The final cytokine concentrations were the following: IL-12 (10ng/ml, R&D Systems), IFN-γ (50ng/ml, R&D Systems) and anti-IFN-γ (10ug/ml, R&D Systems). After 48h of seeding, Treg cells were split into two wells and fresh media without cytokines was added(*45*). Details for reagents and antibodies used are listed in table S1. Sorting strategy is shown in figure S5.

### Single-cell RNA sequencing analyses

Publicly available datasets from LCMV-infected mice (accession number: E-MTAB-8861, EMBL-EBI) and COVID patients (accession number: EGAD00001007718, European Genome-Phenome Archive) were analyzed using R studio (version 2025.05.1+513) and Seurat package (version 5.2.1). Custom R scripts used for the analyses are available from the corresponding author upon request.

### Statistical analyses

Statistical tests were performed using GraphPad Prism (GraphPad Software v10.5.0). Selected statistical tests were indicated in the figure legends. All experiments were performed as 2 to 3 independent repeats, with age– and sex-matched animals across experimental groups. Outliers were identified and excluded using the ROUT test. Relevant significant and non-significant differences were annotated in the figures. Significance was defined as p < 0.05 (*), p < 0.01 (**), p < 0.001 (***), p < 0.0001 (****), or n.s. (not significant, p > 0.05).

## Acknowledgments

We thank the Joller group members for their technical support and helpful discussions. We also thank the group of Prof. Annette Oxenius for providing gp33-monomers and for helpful discussions as well as the team of the Cytometry Facility of the University of Zurich for their support.

## Funding

This work was supported by:

Swiss National Science Foundation grant PP00P3_181037 (NJ)

Swiss National Science Foundation grant 310030_197590 (NJ)

European Research Council grant 677200 Immune Regulation (NJ)

University of Zurich CanDoc fellowship grant FK-24-083 (AEB)

## Author contributions

Conceptualization: AEB, NJ

Investigation: AEB, YMC, LRR

Formal analysis: AEB, YMC

Resources: PZ, SM, BB

Project administration: AEB, NJ

Supervision: NJ

Writing – original draft: AEB, NJ

Writing – review & editing: AEB, YMC, LRR, PZ, SM, BB, NJ

## Competing interests

Authors declare that they have no competing interests.

## Data and materials availability

All data necessary to evaluate the conclusions are presented in the figures or the Supplementary Materials. All scRNA-seq used in this study was previously reported. The mouse data is available in the ArrayExpress database at EMBL-EBI (accession number E-MTAB-8861); the human data is available in the European Genome-Phenome Archive (accession number EGAD00001007718).

## Supplementary Materials

Fig. S1. *Il12rb2* deletion in Treg cells causes impaired type 1 specialization.

Fig. S2. IFN-γ and IL-12 sequentially induce T-bet in Treg cells, without changing their proliferation or viability.

Fig. S3. Tconv cells from Il12rb2 cKO mice are more activated after infection.

Fig. S4. Impaired IL-12 signaling in Treg cells leads to an increase in numbers of IFN-γ^+^ DbGP33^+^ CD8^+^ T cells, without significant differences in viral titers.

Table S1. Reagents and antibodies used.

## Notes

### Competing Interest Statement

The authors have declared no competing interest.

