## Supplementary Materials for "IL-12R signaling promotes type 1 regulatory T cell specialization by sustaining T-bet"

Estrada Brull *et al.*

**This PDF file includes:**

Figs. S1 to S5
Table S1

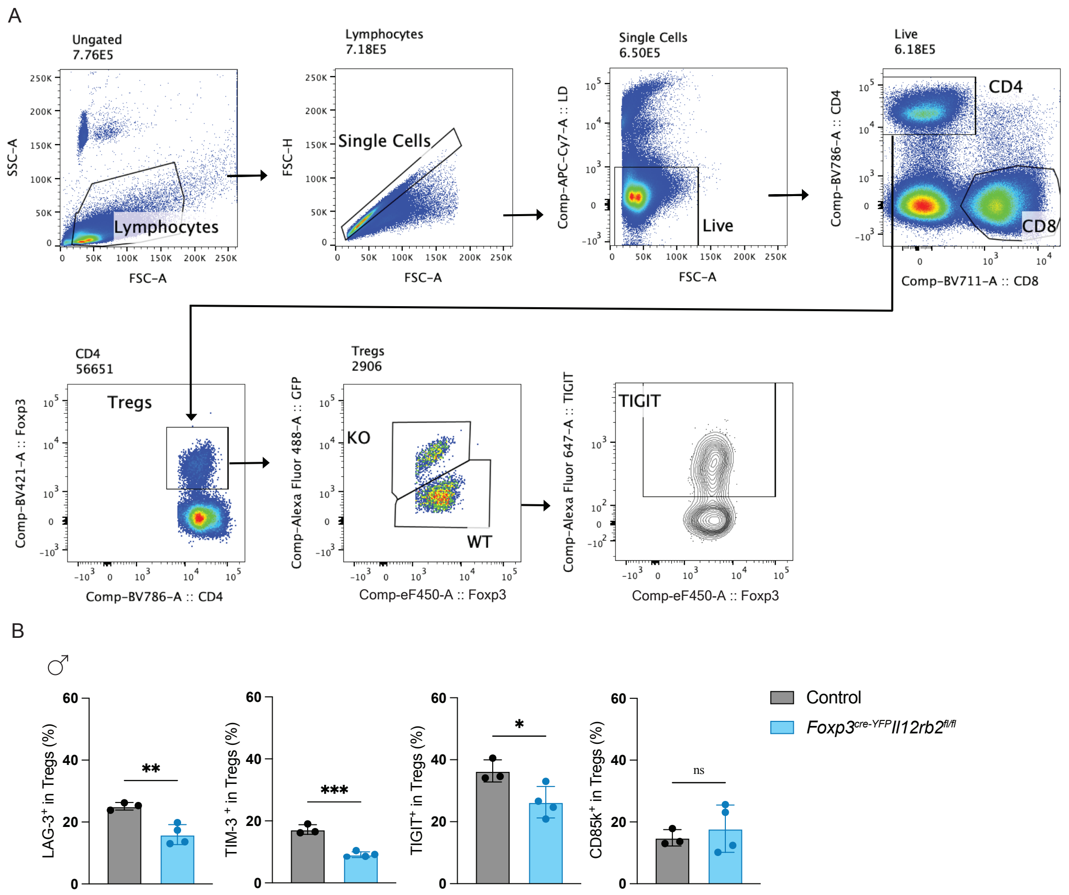

**Fig. S1.** ***Il12rb2*** **deletion in Treg cells causes impaired type 1 specialization.** (**A**) Gating strategy used for flow cytometry analysis. (**B**) Expression of LAG-3, TIM-3, TIGIT and CD85k in *Foxp3$^{cre\text{-}YFP}$Il12rb2$^{fl/fl}$* animals or *Foxp3$^{wt}$Il12rb2$^{fl/fl}$* littermate controls infected with LCMV WE 200 f.f.u at day 10 p.i. Unpaired t tests were used. Data in (B) are from 1 experiment with n=3-4 mice per group. Means with SD are shown. * $p < 0.05$, ** $p < 0.01$, ***$p < 0.001$.

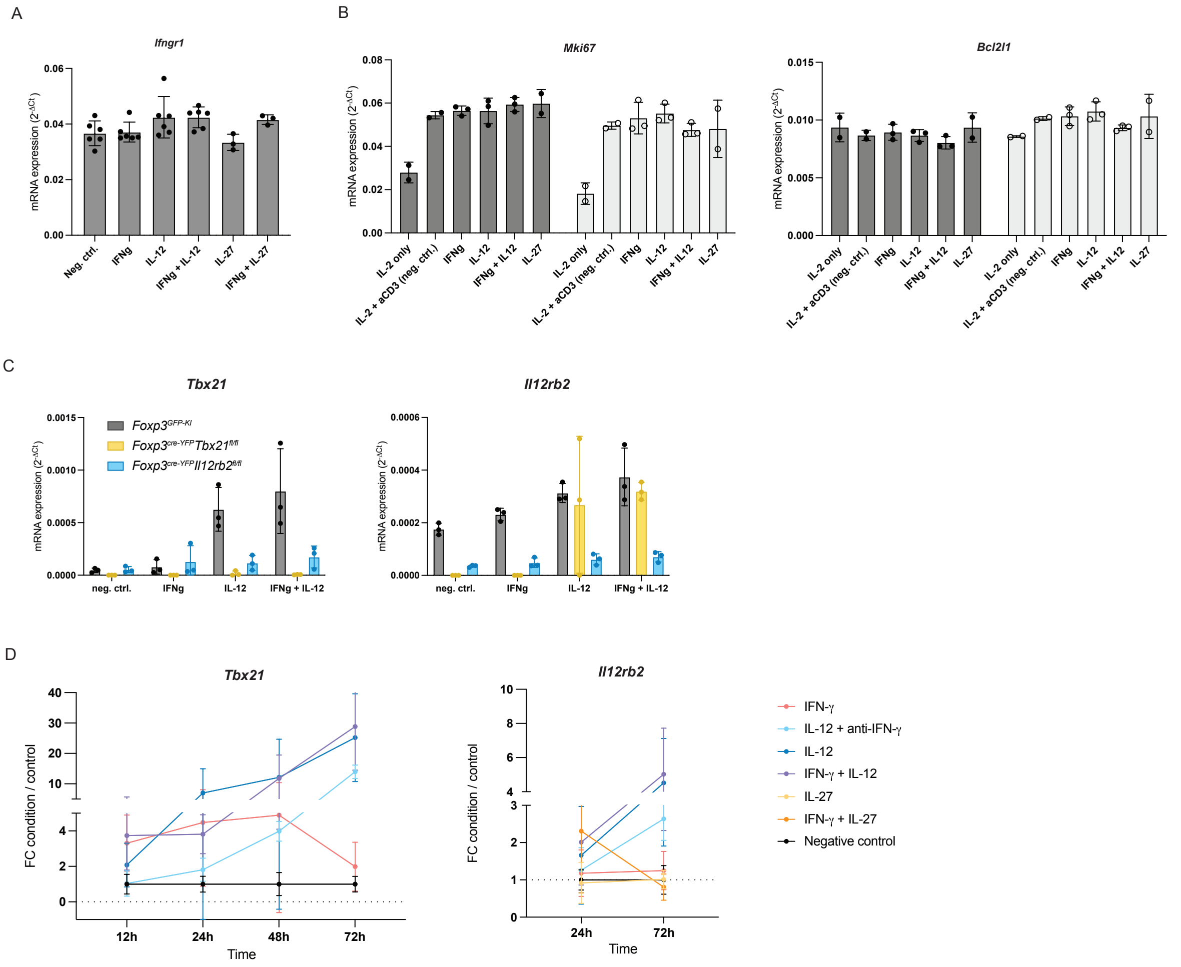

**Fig. S2. IFN-γ and IL-12 sequentially induce T-bet in Treg cells, without changing their proliferation or viability.** (**A**) CXCR3$^-$ Treg cells, or also CXCR3$^+$ Treg cells (**B**), from naive *Foxp3*$^{GFP\text{-}KI}$ were FACS-sorted and incubated with different cytokines for 72h *in vitro* in the presence of IL-2 and anti-CD3 dynabeads. mRNA expression of the specified markers was quantified by RT-qPCR.(**C**) Cxcr3$^-$ Foxp3$^+$ Treg cells from naive *Foxp3*$^{GFP\text{-}KI}$ (WT), *Foxp3*$^{cre\text{-}YFP}$*Tbx21*$^{fl/fl}$ or *Foxp3*$^{cre\text{-}YFP}$*Il12rb2*$^{fl/fl}$ mice were FACS-sorted and incubated with different cytokines for 72h *in vitro* in the presence of IL-2 and anti-CD3 dynabeads. Expression of *Tbx21* and *Il12rb2* mRNA was quantified. (**D**) CXCR3$^-$ Treg cells from naive *Foxp3*$^{GFP\text{-}KI}$ mice were FACS-sorted and incubated with different cytokines for 12h, 24h, 48h or 72h *in vitro* in the presence of IL-2 and anti-CD3 dynabeads. Plot indicating the fold change of *Tbx21* mRNA (dCT cytokine condition divided by dCT of the negative control) expression over time is shown. Data in (A-C) are pooled from 1-2 independent experiments with n=2-6 technical replicates per group. Data in (D) are calculated from 2-5 independent experiments with n=3-15 technical replicates per group. Means with SD are shown.

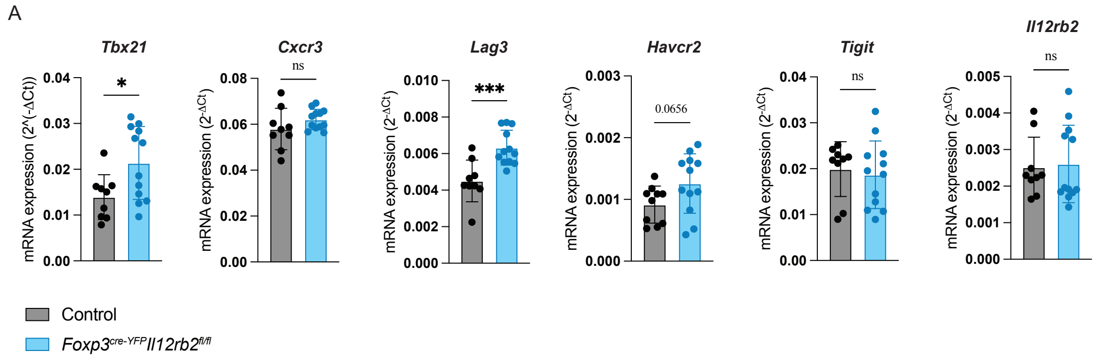

**Fig. S3. Tconv cells from Il12rb2 cKO mice are more activated after infection.** (**A**) mRNA expression in CD44$^{hi}$ Tconv cells isolated from *Foxp3$^{GFP\text{-}KI}$* or *Foxp3$^{cre\text{-}YFP}$Il12rb2$^{fl/fl}$* mice at day 10 after infection with LCMV WE 200 f.f.u. Data are pooled from 4 independent sorts with n=9-12 mice per group. Unpaired t tests were used. Means with SD are shown. * $p < 0.05$, *** $p < 0.001$.

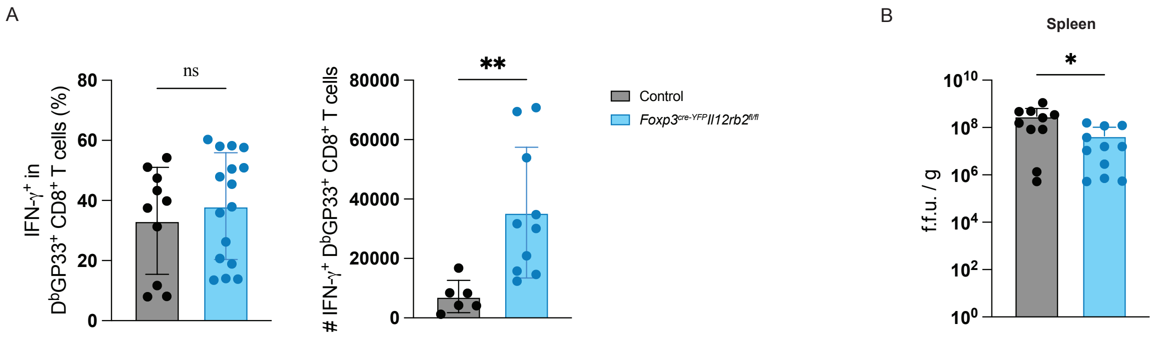

**Fig. S4. Impaired IL-12R signaling in Treg cells leads to an increase in numbers of IFN-γ$^{+}$ DbGP33$^{+}$ CD8$^{+}$ T cells, with differences in viral titers.** (**A**) Percentage and total numbers in spleen of IFN-γ$^{+}$ DbGP33$^{+}$ CD8$^{+}$ T cells in *Foxp3$^{cre\text{-}YFP}$* or *Foxp3$^{cre\text{-}YFP}$Il12rb2$^{fl/fl}$* mice at day 27 after infection with LCMV Cl13 2x10$^{6}$ f.f.u. (**B**) Viral titers in spleens from *Foxp3$^{cre\text{-}YFP}$* or *Foxp3$^{cre\text{-}YFP}$Il12rb2$^{fl/fl}$* mice at day 27 after infection with LCMV Cl13 2x10$^{6}$ f.f.u. measured by RT-qPCR. Data are pooled from 2-3 independent experiments with n=6-16 animals per group. Means with SD are shown. Unpaired t tests were used. ** $p < 0.01$.

A

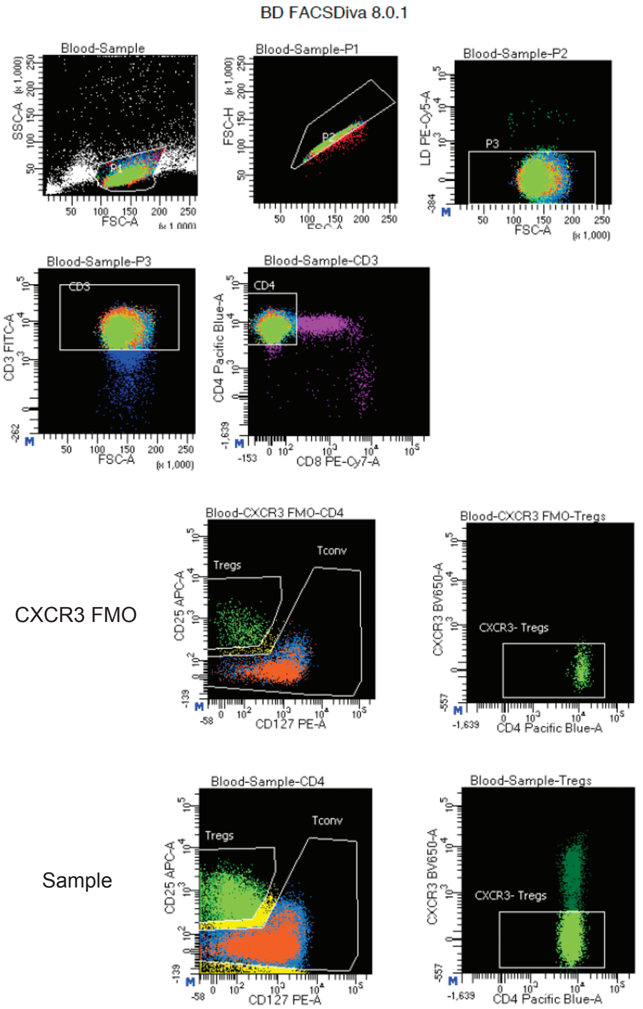

**Fig. S5. Sorting of CXCR3$^-$ human Treg cells.** (**A**) Sorting strategy for isolating CD4$^+$CD127$^{low}$CD25$^+$CXCR3$^-$ Treg cells from PBMCs of healthy donors.

| Media and common reagents | | |
|---|---|---|
| **Name** | **Order no.** | **Company** |
| **RPMI 1640 medium** | 21875034 | Gibco |
| **Minimum Essential Medium** | 21090-055 | Gibco |
| **Penicillin-Streptomycin** | 15140-122 | Gibco |
| **Glutamine** | 25030-024 | Gibco |
| **FBS** | 35-079-CV | Corning |
| **FBS** | Lot CP22-5296 | Capricorn |
| **Brefeldin A** | 420601 | BioLegend |
| **gp33 (KAVYNFATM)** | custom made | EMC microcollections |
| **gp61 (GLKGPDIYKGVYQFKSVEFD)** | custom made | BioCat |
| **aCD3** | #BE0001-1 | BioXcell |
| **β-mercaptoethanol** | A1108,0100 | PanReac AppliChem |
| **Sodium Pyruvate** | 11360039 | Gibco |
| **Non-Essential aminoacids** | 11140035 | Gibco |
| **MEM vitamins** | 11120037 | Gibco |
| **HEPES** | 15630056 | Gibco |

| Lung processing | | |
|---|---|---|
| **Name** | **Order no.** | **Company** |
| **Collagenase I** | 11088866001 | Roche |
| **DNAse I** | 60852700 | Roche |
| **MACS C-tubes** | 130-093-237 | Milteny Biotec |
| **Percoll** | 17-0891-01 | GE Healthcare |

| Cell isolation and magnetic sorting | | |
|---|---|---|
| **Reagents** | | |
| **Name** | **Order no.** | **Company** |

| | | |
|---|---|---|
| **MojoSort Mouse CD4 Nanobeads** | 480070 | BioLegend |
| **MojoSort Mouse CD4 T Cell Isolation Kit** | 480033 | BioLegend |
| **MojoSort Mouse CD8 T cell Isolation kit** | 480035 | BioLegend |
| **CD4 MicroBeads (human)** | 130-045-101 | Miltenyi |
| **CD14 MicroBeads (human)** | 130-050-201 | Miltenyi |
| **7-AAD Viability Staining Solution** | 420404 | BioLegend |
| **Human TruStain FcX™ (Fc Receptor Blocking Solution)** | 422301 | BioLegend |
| **Flow cytometry** | | |
| **Buffers and reagents** | | |
| **Name** | **Order no.** | **Company** |
| **Foxp3 / Transcription Factor Fixation/Permeabilization Concentrate and Diluent** | 00-5521-00 | Invitrogen (ThermoFischer) |
| **BD Cytofix/Cytoperm** | 554722 | BD Biosciences |
| **Permeabilization Buffer (10X)** | 00-8333-56 | eBiosciences (ThermoFisher) |
| **UltraComp eBeads Compensation Beads, 5ml** | 01-2222-42 | Invitrogen |
| **CountBright Absolute Counting beads** | C36950 | invitrogen |
| **CellTrace Violet Cell Proliferation Kit** | C34571 | ThermoFischer |
| **RNA extraction, RT & RT-PCR** | | |
| **Name** | **Order no.** | **Company** |
| **Rneasy minikit** | 74106 | Qiagen |
| **QIAamp Viral RNA Mini Kit** | 52906 | Qiagen |
| **High-Capacity cDNA RT kit** | 4368814 | Life Technologies |

| | | |
|---|---|---|
| **TaqMan Fast Advanced Master Mix** | 4444963 | Applied Biosystems |
| **Probes mouse** | | |
| **actb** | 4352341E | Applied Biosystems |
| **Cxcr3** | Mm99999054_s1 | Applied Biosystems |
| **Foxp3** | Mm00475162_m1 | Applied Biosystems |
| **Havcr2** | Mm00454540_m1 | Applied Biosystems |
| **Ifng** | Mm01168134_m1 | Applied Biosystems |
| **Ifngr1** | Mm00599890_m1 | Applied Biosystems |
| **Il12rb2** | Mm00434200_m1 | Applied Biosystems |
| **Lag3** | Mm00493071_m1 | Applied Biosystems |
| **Lilrb4** | Mm01614371_m1 | Applied Biosystems |
| **Tbx21** | Mm00450960_m1 | Applied Biosystems |
| **TIGIT Forward primer: 5′-CTGATACAGGCTGCCTTCCT-3′** | | Sigma-Aldrich |
| **TIGIT Reverse primer: 5′-TGGGTCACTTCAGCTGTGTC-3′** | | Sigma-Aldrich |
| **TIGIT Probe: 5′-AGGAGCCACAGCAGGCACGA-3′ (FAM, TAMRA)** | | Sigma-Aldrich |
| **LCMV Forward primer: 5′-CTGACGAGGTCAACCCGG-3′** | | Sigma-Aldrich |
| **LCMV Reverse primer: 5′-CAAGTACTCACACGGCATGGA -3′** | | Sigma-Aldrich |
| **LCMV Probe: 5′-CTTGCCGACCTCTTCAATGCGCAA -3′ (FAM, TAMRA)** | | Sigma-Aldrich |
| **Probes human** | | |
| **ACTB** | Hs99999903_m1 | Applied Biosystems |

| | | |
|---|---|---|
| **CXCR3** | Hs00171041_m1 | Applied Biosystems |
| **FOXP3** | Hs01085834_m1 | Applied Biosystems |
| **IFNGR1** | Hs00988304_m1 | Applied Biosystems |
| **IL12RB2** | Hs00155486_m1 | Applied Biosystems |
| **TBX21** | Hs00894392_m1 | Applied Biosystems |

| ***In vitro* Treg assays** | | |
|---|---|---|
| **Mouse** | | |
| **Name** | **Order no.** | **Company** |
| **Dynabeads™ Mouse T-Activator CD3/CD28 for T-Cell Expansion and Activation** | 11452D | ThermoFischer |
| **Recombinant mouse IL-2** | 575402 | BioLegend |
| **Recombinant Mouse IL-12 (p70)** | 577006 | BioLegend |
| **Recombinant Mouse IL-27** | 577404 | BioLegend |
| **Recombinant mouse IFN-γ** | 714006 | BioLegend |
| **LEAF Purified anit-mouse IFN-γ (Clone: XMG1.2)** | 505827 | BioLegend |

| **Human** | | |
|---|---|---|
| **Name** | **Order no.** | **Company** |
| **Dynabeads™ Human T-Activator CD3/CD28 for T Cell Expansion and Activation** | 11161D | Gibco |
| **Recombinant human IL-12, 5ug** | 219-IL-005 | R&D Systems |
| **Recombinant human IL-2, 10ug** | 202-IL-010 | R&D Systems |
| **Recombinant human IL-7, 5ug** | 207-IL-005 | R&D Systems |
| **Recombinant human IFNg, 100ug** | 285-IF-100 | R&D Systems |
| **Human IFN-gamma Antibody, 25ug** | MAB285-SP | R&D Systems |

| Blood collection and PBMC isolation | | |
|---|---|---|
| **Name** | **Order no.** | **Company** |
| **EDTA Vacutainer tubes** | 367525 | BD Biosciences |
| **Ficoll Paque** | 17544202 | Cytiva |

| Antibodies | | | | |
|---|---|---|---|---|
| **Cell sorting - mouse** | | | | |
| **Marker (clone)** | **Fluorochrome** | **Order no.** | **Company** | **Dilution** |
| **CD4 (GK1.5)** | APC | 100412 | BioLegend | 1:200 |
| **CD44 (IM7)** | AF700 | 103026 | BioLegend | 1:300 |
| **CXCR3 (CXCR3-173)** | PE | 126506 | BioLegend | 1:100 |
| **CD25 (3C7)** | PE | 101904 | BioLegend | 1:100 |

| Cell sorting - human | | | | |
|---|---|---|---|---|
| **Marker (clone)** | **Fluorochrome** | **Order no.** | **Company** | **Dilution** |
| **CD3 (OKT3)** | FITC | 317306 | BioLegend | 1:200 |
| **CD4 (SK3)** | BUV496 | 612937 | BD | 1:200 |
| **CD4 (SK3)** | Pacific Blue | 344619 | BioLegend | 1:400 |
| **CD25 (M-A251)** | APC | 356110 | BioLegend | 1:50 |
| **CD45RA (HI100)** | BV510 | 304141 | BioLegend | 1:200 |
| **CD127 (A019D5)** | PE | 351304 | BioLegend | 1:100 |
| **CXCR3 (G025H7)** | BV650 | 353729 | BioLegend | 1:100 |

| **Flow cytometry acquisition** | | | | |
|---|---|---|---|---|
| **Marker (clone)** | **Fluorochrome** | **Order no.** | **Company** | **Dilution** |
| **LIVE/DEAD™ Fixable Blue Dead Cell Stain Kit** | | L-34962 | Invitrogen | 1:500 |
| **Zombie NIR Fixable Viability Kit** | | 423106 | BioLegend | 1:500 |
| **CD25 (PC61)** | Biotin | 102003 | BioLegend | 1:500 |
| **CD25 (PC61)** | BV650 | 102038 | BioLegend | 1:300 |
| **CD4 (RM4-5)** | BUV496 | 741050 | BD | 1:500 |
| **CD4 (RM4-5)** | BV421 | 100544 | BioLegend | 1:200 |
| **CD4 (RM4-5)** | BV785 | 100552 | BioLegend | 1:200 |
| **CD4 (RM4-5)** | PerCP-Cy5.5 | 100540 | BioLegend | 1:200 |
| **CD44 (IM7)** | APC-Fire750 | 103061 | BioLegend | 1:300 |
| **CD44 (IM7)** | PE-Cy5 | 103009 | BioLegend | 1:400 |
| **CD44 (IM7)** | PerCP | 103036 | BioLegend | 1:400 |
| **CD45.1 (A20)** | BUV805 | 741958 | BD | 1:200 |
| **CD8 (53-6.7)** | BUV395 | 565968 | BD | 1:500 |
| **CD8 (53-6.7)** | BV711 | 100748 | BioLegend | 1:500 |
| **CD85K (H1.1)** | AF647 | 144906 | BioLegend | 1:100 |
| **CX3CR1 (SA011F11)** | Pacific Blue | 149037 | BioLegend | 1:400 |
| **CXCR3 (CXCR3-173)** | Biotin | 126503 | BioLegend | 1:100 |
| **CXCR3 (CXCR3-173)** | BV510 | 126528 | BioLegend | 1:200 |
| **CXCR3 (CXCR3-173)** | BV650 | 126531 | BioLegend | 1:100 |
| **CXCR3 (CXCR3-173)** | PE | 126505 | BioLegend | 1:100 |
| **CXCR6 (SA051D1)** | APC-Fire750 | 151130 | BioLegend | 1:50 |
| **FOXP3 (FJK-16s)** | eF450 | 48-5773-82 | eBioscience | 1:200 |
| **FOXP3 (FJK-16s)** | FITC | 11-5773-82 | eBioscience | 1:200 |
| **GFP/YFP-TAG (FM264G)** | AF488 | 338008 | BioLegend | 1:400 |
| **IFNg (XMG1.2)** | APC | 505810 | BioLegend | 1:300 |

| | | | | |
|---|---|---|---|---|
| **IFNg (XMG1.2)** | PE | 505808 | BioLegend | 1:300 |
| **IFNg (XMG1.2)** | PE-Dazzle594 | 505845 | BioLegend | 1:500 |
| **KLRG1 (2F1)** | SB702 | 67-5893-82 | eBioscience | 1:100 |
| **LAG3 (C9B7W)** | Biotin | 125206 | BioLegend | 1:100 |
| **LAG3 (C9B7W)** | BV421 | 125221 | BioLegend | 1:100 |
| **LAG3 (C9B7W)** | BV785 | 125219 | BioLegend | 1:50 |
| **Ly108 (330-AJ)** | PE | 134605 | BioLegend | 1:200 |
| **NRP1 (V46-1954)** | BV711 | 752456 | BD | 1:500 |
| **PD-1 (29F.1A12)** | BV605 | 135220 | BioLegend | 1:100 |
| **PD-1 (RMP1-30)** | BUV737 | 568363 | BD | 1:100 |
| **TBET (4B10)** | PE/Cy7 | 644824 | BioLegend | 1:300 |
| **TCF1/7 (S33-966)** | R718 | 567587 | BD | 1:100 |
| **TIGIT (1G9)** | BV421 | 142111 | BioLegend | 1:50 |
| **TIGIT (1G9)** | PE-Dazzle594 | 142110 | BioLegend | 1:50 |
| **TIM3 (5D12/TIM3)** | BB700 | 747619 | BD | 1:200 |
| **TIM3 (RMT3-23)** | Biotin | 119720 | BioLegend | 1:100 |
| **TNFa (MP6-XT22)** | PE/Cy7 | 506324 | BioLegend | 1:500 |
| **TOX (TXRX10)** | PE | 12-6502-80 | Invitrogen | 1:300 |
| **Streptavidin** | APC | 405243 | BioLegend | 1:500 |
| **Streptavidin** | BV480 | 564876 | BD | 1:500 |
| **Streptavidin** | PE-Cy5 | 405205 | BioLegend | 1:400 |
| **In vivo Antibodies** | | | | |
| **Name** | **Clone** | **Order no.** | **Company** | |
| **InVivoMAb anti-mouse IL-27 p28** | MM27.7B1 | BE0326-5mg-A | BioXCell | |
| **InVivoMAb mouse IgG2a isotype control** | C1.18.4 | BE0085-5mg-A | BioXCell | |

**Table S1. Reagents and antibodies used.**
